# Assessing climate change risks associated with precipitation at unfavorable times in winter wheat using an improved crop calendar model incorporating vernalization and winter survival

**DOI:** 10.1101/2023.10.19.563190

**Authors:** Keach Murakami, Toshichika Iizumi, Seiji Shimoda

## Abstract

Crop phenology calendars are necessary for designing breeding goals and for developing effective management practices. Winter wheat is a representative biennial, the cultivation schedule of which is constrained by winter climate conditions, particularly the processes of vernalization and winter survival. Here, we present improvements to a rule-based crop phenology model by incorporating these factors so that it can be used to accurately estimate the phenological events of winter wheat from daily meteorological data. We tested the improved model in Hokkaido, the northernmost Japanese island, which is characterized by seasonal snow cover and a wet summer. The results confirmed that implementing these factors contributed to accurate estimates of peak occurrence dates of winter wheat phenological events. Furthermore, we applied the improved model to simulate wheat phenology under 2 K and 4 K warmer scenarios. The results showed a delayed sowing period up to approximately one month and slight advancements in both flowering and harvesting, leading to a shorter growth period. While this shortened period may be largely compensated by a decrease in the snow-covered period, the shifts in the vegetative and reproductive phases may have a significant influence on sink-source balance of wheat. We also assessed the risks of pollination failure and preharvest sprouting, both of which are associated with the timing of precipitation, based on the number of rainy days around flowering, and the incidence of precipitation over two consecutive days around the time of harvesting. Our simulations suggested increased risk of pollination failure and reduced risk of preharvest sprouting, leading to an increase in the probability of crop failure. These findings underscore the importance of implementing adaptive measures to mitigate precipitation-related risk under future climate scenarios. Further, the findings provide valuable insights for winter wheat breeders and agronomists, thereby facilitating crop production adaptation strategies.

## Introduction

The adjustment of crop calendars in response to climate change serve as the basis for formulating breeding goals and management practices aimed at adaptation (e.g., plant density, fertilization, and pest and disease control). Among major crops, the winter wheat calendars for regions in the mid- and high latitudes, which are characterized by extensive snow covers and low temperatures, would shift markedly in a warmer climate. In addition to changes driven by rising temperatures, some of the development stages of winter wheat are susceptible to wet conditions. For example, shifts in wheat phenology in conjunction with changes in summer precipitation patterns could decrease grain productivity and quality. Pollination failure due to precipitation at the time of flowering (Shimoda *et al* 2022, Nóia Júnior *et al* 2023) and preharvest sprouting as a result of precipitation around the harvesting (Mares and Mrva 2014) are both known risk factors for winter wheat production associated with precipitation. Although these risks have attracted less attention in major wheat producing areas with dry climates, and since no widely used model or index has been developed to assess these risks, they have been a major concern in wheat production in humid environments, such as Japan. In Hokkaido, the northernmost Japanese island, winter wheat is widely cultivated under snow cover from winter to spring and wet climate in summer. The humid conditions have historically caused significant yield reductions in this region (Shimoda *et al* 2022). Furthermore, because climate models have suggested an increase in summer precipitation in this region (Takabatake and Inatsu 2022), risks associated with summer precipitation should be seriously considered to ensure wheat production in this region. Climate projections have suggested intensified precipitation worldwide (Masson-Delmotte *et al* 2021), implying that there is an increased likelihood for wheat farmers to be adversely affected by precipitation occurring at unfavorable times. Therefore, assessments of precipitation-induced risks that combine future changes in crop calendars and precipitation patterns are important for developing optimal adaptation strategies.

Numerous studies on crop calendars have developed datasets and models. Some of these models are driven by meteorological data and are well suited for estimating crop calendars in a changing climate. For a simple example, a model developed by the World Meteorological Organization (WMO) can be used to determine sowing and harvesting windows based on whether the climatological duration of the rainy season is sufficient to complete a crop cycle (de Roos 2021). Waha *et al* (2012) developed a model that estimates sowing months for 11 major crops based on the intra-annual variability in monthly temperatures and precipitations. Minoli *et al* (2019) proposed an algorithm that calculates the maturity date of crops based on the sowing date and monthly climate data using species-specific vegetative and reproductive durations. Further, Minoli *et al* (2022) also estimated growth periods of several crops at the end of the century by projecting these calendars models (Waha *et al* 2012, Minoli *et al* 2019) and simulated the crop yields using a biophysical crop model. Iizumi *et al* (2019) proposed a rule-based agro-climatic resource-based crop calendar model forced by 10-year daily meteorological data that can be used to estimate suitable periods (i.e. windows) for sowing and harvesting. Specifically, their model considers crop-specific biological requirements related to heat, chilling, and moisture, as well as the extent to which field workability is restricted by snow covers and intense precipitation. While the estimated sowing and harvesting windows for summer crops are in close agreement with crop progress reports, there is scope for model improvements for winter wheat, particularly to address the issue of overly extended estimated windows. The shortfall of estimated calendar is partly due to the inadequate representation of the vernalization process in the model. Vernalization refers to a physiological process in which a certain amount of chilling (i.e., low temperatures) is required to initiate panicle formation in biennial plants such as winter wheat (Amasino 2004). When winter wheat is sown in spring or early summer, plants do not produce any grains in spite of sufficient thermal resources. Another process not considered in the model of Iizumi *et al* (2019) is winter survival. To ensure successful winter survival in regions that experience intensive seasonal snowfall, wheat seedlings must reach a certain size before snow cover. Late sowing results in there being only a small carbohydrate reserve before snow cover, leading to lower resistance to mold under the snowpack (Bruehl and Cunfer 1971, Mohammad *et al* 1997, Yoshida *et al* 1998). Insufficient root growth in seedlings makes them prone to winter damage, mainly due to limited water uptake caused by low soil temperatures and physical injury induced by frost heaving. Since vernalization and winter survival are key factors affecting the design of winter wheat calendar, these processes need to be considered in crop calendar models for accurate estimation of winter wheat calendars in a changing climate.

This study aimed to 1) improve the crop calendar model proposed by Iizumi *et al* (2019) to more accurately estimate winter wheat calendars by incorporating vernalization and winter survival; 2) apply the improved model to project shifts in winter wheat calendar under projected climates; and 3) assess the risks of pollination failure and preharvest sprouting and the resulting opportunity loss of winter wheat cultivation under compound changes in the calendar and in precipitation patterns. We focused on Hokkaido (figure 1), a good testbed for the model improvement and for studying precipitation-induced risks associated with winter wheat production.

**Figure 1:**
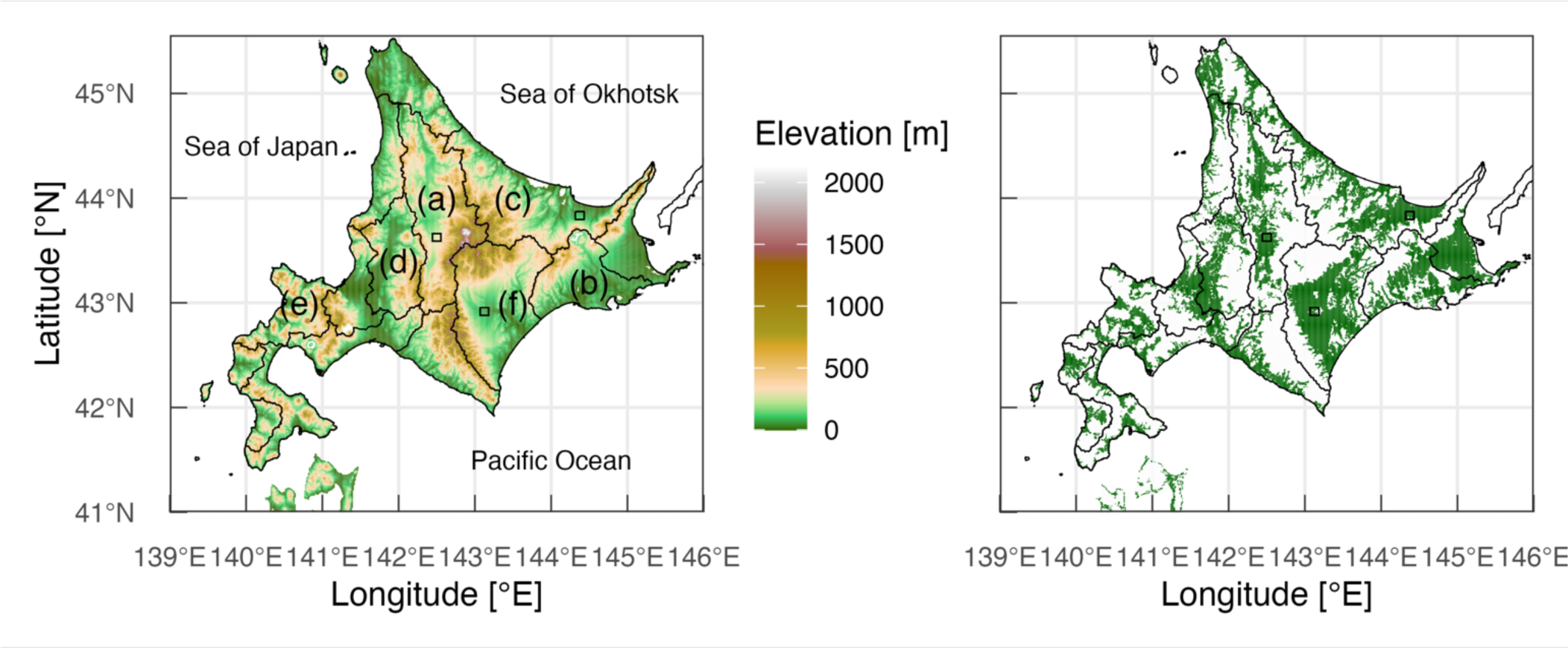
Locations of study sites (left) and the distribution of upland crop fields (shown in green; right) in Hokkaido. Three squares indicate the sites subjected to the risk assessment of precipitation-related crop failure. The black boundaries denote the subprefectural administrative units: a, Kamikawa; b, Kushiro; c, Okhotsk; d, Sorachi; e, Shiribeshi; and f, Tokachi. The boundaries and land-use data are from the National Land Information Division, National Spatial Planning and Regional Policy Bureau (https://nlftp.mlit.go.jp/ksj/index.html, accessed on 2023– 09–15).

## Materials and Methods

### 1.1 Crop calendar model

The crop calendar model originally proposed by Iizumi *et al* (2019) and modified in this study estimates likely time windows for sowing and harvesting at a daily resolution by considering constraints such as crop’s biological requirements for heat, chilling and water, as well as field workability, which is affected by snow cover and intense precipitation around sowing and harvesting.

We augmented the original model with two rules: one related to vernalization and another to winter survival. Then, we added rules regarding pollination failure and preharvest sprouting to assess the opportunity loss due to precipitation-associated crop failure (see ‘Rain risk assessment’ for details). The low temperature requirement for vernalization was incorporated into the model instead of the chilling requirement in the original model. In the improved model, it was assumed that vernalization is completed after 65 cold days (daily mean temperature ≤ 5 °C) from sowing; this criterion is sufficient to induce full vernalization in various cultivars (Kato and Yamagata 1988). Wheat plants were assumed not to produce grains if vernalization had not been completed by the panicle formation stage. Panicle formation corresponded to a cumulative effective temperature of 700 °C d after sowing with a base temperature (T_base_) of 0 °C. This value was determined based on ground-truth phenology data and daily meteorological data summarized for each subprefecture (detailed in the following sections).

Another rule on winter survival was set based on the growing-degree days (GDD) after sowing to the commencement of snow cover. In this study, we defined snow cover as the period when the maximum number of consecutive days with a non-zero snow water equivalent. If that period was shorter than 30 d, then this rule was skipped in the model. Local guidelines for farmers in the study area suggest that the required GDD for winter survival ranges from 430 to 640 °C d (T_base_ = 3 °C) from sowing to November 15, depending on cultivars and subprefectural regions (Chiba 2023). However, we did not adopt this threshold because actual sowing dates were later than those calculated using the local guidelines in most regions and years, and we adopted a GDD of 300 °C d and a T_base_ of 0 °C instead (see also figure S1).

In addition to the windows for sowing and harvesting, the model also estimate time windows for emergence and flowering according to Mori *et al* (2023); plants were assumed to reach these phenological stages when the GDD after sowing reached 6% and 59% of the crop total thermal requirement. Since the present analyses focused on Hokkaido, a region with abundant spring snowmelt and a rainy summer, we did not consider the limitation of water availability during the crop duration. Available evidence indicates that summer precipitation in this region is likely to increase (Takabatake and Inatsu 2022), suggesting that the water availability would not be a limiting factor.

### 1.2 Phenology data

Ground truth data for the occurrences of winter wheat phenological events (sowing, germination, panicle formation, and harvesting) from 2000 to 2020 were collected from crop progress reports published by the agricultural administration department of the Hokkaido prefectural government (https://www.pref.hokkaido.lg.jp/ns/gjf/seiiku/). We used data from six subprefectures where data for more than four years were available (a–f in figure 1).

### 1.3 Meteorological datasets

In this study, we used two meteorological datasets compiled in the Agro-Meteorological Grid Square Data System (Ohno *et al* 2016) operated by the National Agriculture and Food Research Organization (NARO): a historical dataset spanning from 2000 to 2020 (HIST) and a dataset of future projections from 2070 to 2090. The former was generated by interpolating data observed at weather stations operated by the Japan Meteorological Agency. The latter was generated by applying bias correction and statistical downscaling to outputs of five global climate models (CSIRO-Mk3.6.0, GFDL-CM3, HadGEM2-ES, MIROC5, and MRI-CGCM3) listed in the Coupled Model Intercomparison Project Phase 5 (CMIP5). Two representative concentration pathways (RCP2.6 and RCP8.5) used here represent global temperature increase below 2 K by and approximately 4 K by around 2100, relative to pre-industrial levels, respectively. From these datasets, we retrieved the 1-km resolution daily meteorological data—mean, maximum, and minimum air temperatures and precipitation—using the interface for this system (Murakami and Nagasaki 2023) within the R software (R Core Team 2022).

Although information on snow cover is necessary to calculate field workability, it is not available in the datasets. We therefore estimated the snow water equivalent from daily temperature and precipitation data according to Trnka *et al* (2010). In this study, a snow water equivalent value greater than zero indicated the presence of snow cover.

For analyses at the subprefecture scale, meteorological data were aggregated at the same scale to maintain spatial consistency. All grid cells with upland crop fields were extracted and their mean values were calculated. Land use data (as of 2009) generated by the Ministry of Land, Infrastructure, Transport and Tourism (https://nlftp.mlit.go.jp/ksj/gml/datalist/KsjTmplt-L03-b.html) were employed to identify grid cells with upland crop fields.

### 1.4 Rain risk assessment

Focusing on the flowering and harvesting periods in which wheat is susceptible to precipitation events, we assessed the risks of pollination failure and preharvest sprouting. The risk of pollination failure was quantified as the number of rainy days within ±5 d of the peak date of the flowering window. A day with precipitation >1 mm d–1 was defined as a rainy day. For preharvest sprouting, we used the definitions of Mares and Mrva (2014), who stated that ‘around 10–15 mm of rain is probably the minimum required’ and ‘the grain will need to stay moist for around 2 days’ to initiate sprouting. The risk of preharvest spouting was quantified as the maximum amount of consecutive 2-day precipitation within ±5 d of the peak date of the harvesting window. These risks were calculated at the subprefecture level.

We also calculated phenological event windows by imposing additional rules to avoid unfavorable precipitation events during the flowering and harvesting periods. The estimated daily likelihood value of sowing indicates the chance of completing an annual crop lifecycle in a given 20-year period when the crop is planted on a given day. Daily sowing likelihoods calculated with and without these rules were integrated over time to evaluate the precipitation-associated opportunity loss of winter wheat cultivation under historical and future climates (figure S2). Pollination failure and preharvest sprouting were assumed to occur when there were six or more rainy days (i.e. more than half) and two or more consecutive days with daily precipitation of 15 mm or greater within ±5 d of the event dates, respectively. The integrated sowing likelihoods were compared among three small areas (∼ 100 km2) (square insets in figure 1).

### 1.5 Statistical analyses

A two-way analysis of variance (ANOVA) was applied to assess temporal and spatial effects on rain-associated risks. The effects of climate scenarios (HIST, RCP2.6, and RCP8.5), and subprefectures (fourteen regions), and their interactions were analyzed. The significance of the differences among the mean values under three climate scenarios were tested by Tukey–Kramer’s HSD test. All analyses were performed using the R software (R Core Team 2022).

## Results and Discussion

### 2.1 Model improvements

When compared to the original model, the peak dates of the phenological event windows estimated using the improved model were in close accordance with the ground truth data (figure 2). Most of the peak dates estimated by the improved model fell within the reported ranges. Notably, the improved model accurately estimated germination in early October, a feature that the original model was unable to capture (Iizumi *et al* 2019, Mori *et al* 2023). Although flowering dates were not available in the ground truth dataset, the estimated flowering dates (middle of June) were also considered reasonable in light of our field experience. These results indicate the validity of the improved model for the study area. In addition, the improved model showed narrower windows than the original model (Iizumi *et al* 2019, Mori *et al* 2023)(figure 3). This was because the additional rules representing the vernalization and winter survival had eliminated unreasonable early and late sowing in the original model. These modifications worked well for the inland area with intensive snowfall (a-Kamikawa in figure 1) and plain area facing the Pacific Ocean characterized by a drier winter (f-Tokachi in figure 1). As a result of the narrower sowing window, the windows of the other phenological events were also narrowed. Narrowing phenological windows might be particularly useful when one considers multiple crops are cultivated in a year, which plays a key role in gaining crop production under the climate change (Iizumi and Ramankutty 2015, Urfels *et al* 2022).

**Figure 2:**
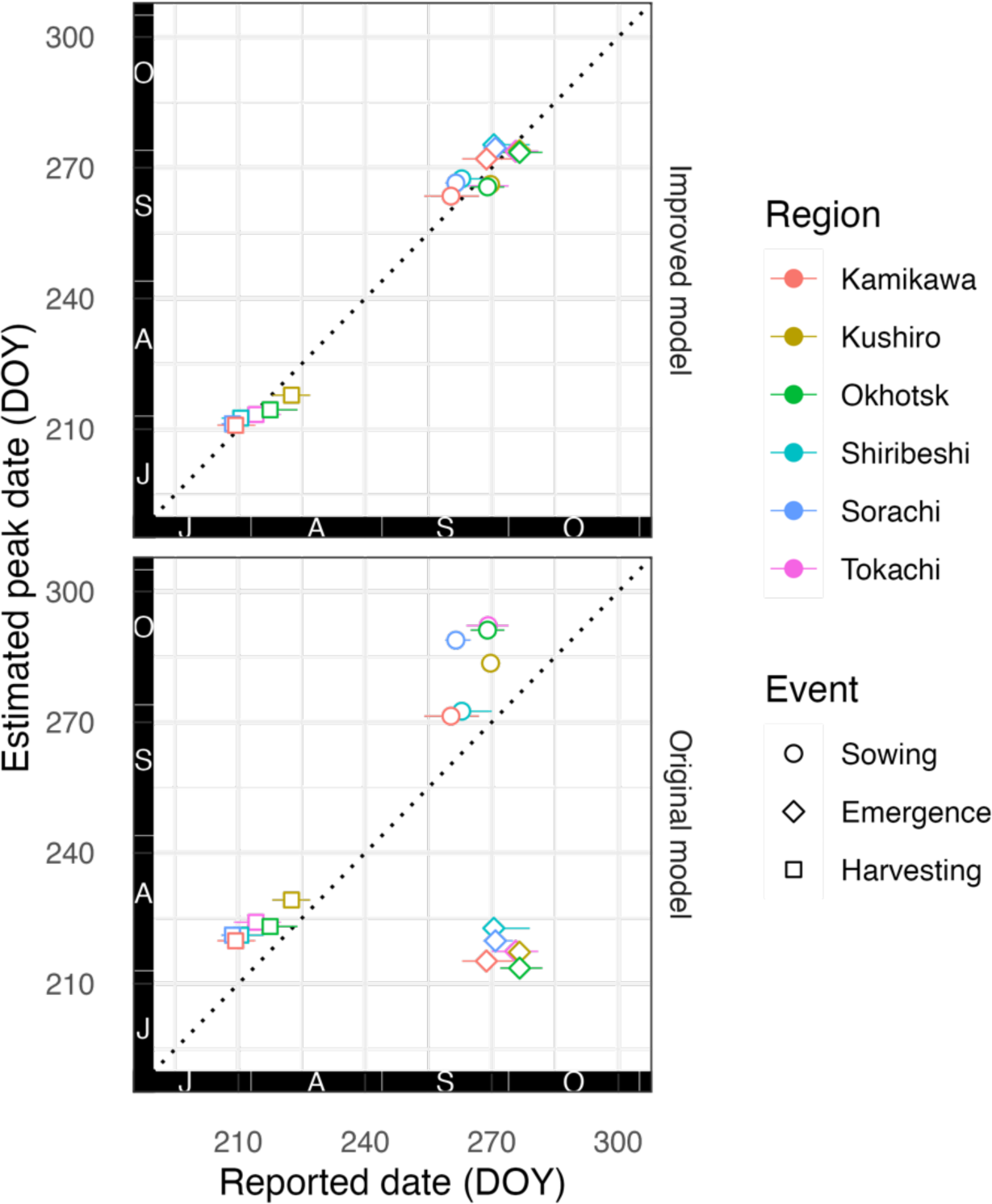
Comparisons between the reported and estimated dates of winter wheat phenological events. Horizontal bars represent the reported minimum to maximum dates in day of the year (DOY) (*N* = 5–21).

**Figure 3:**
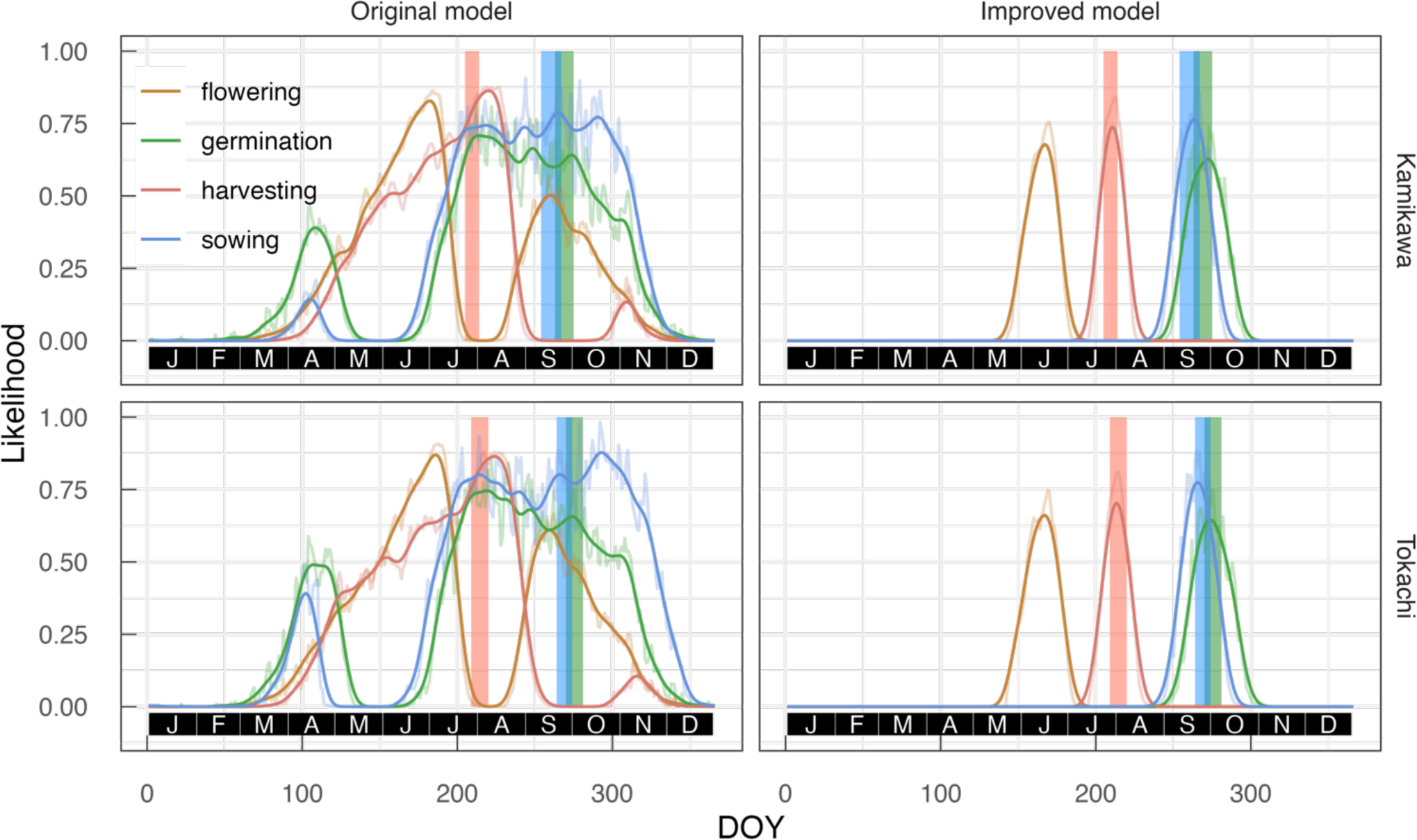
Likelihood of the occurrences of winter wheat phenological events estimated by the original and improved models. Shaded bars indicate the reported ranges of phenological events (2000–2020; *N* = 6–21). For the peaks, thin and thick lines indicate the raw model output and smoothed data (5-day moving average repeated 10 times), respectively. DOY: Day of the year.

### 2.2 Projected calendar shifts

Projected temperature increases would delay sowing by 7–11 d under the RCP2.6 scenario and by 21–26 d under the RCP8.5 scenario by the end of the 21st century (figures 4 and S3). These changes were caused by warming and a delayed commencement of snow cover. Flowering was projected to move forward by 1–3 d under the RCP2.6 scenario and 6–10 d under the RCP8.5 scenario and harvesting was by approximately 0–2 d under the RCP2.6 scenario and 3–5 d under the RCP8.5 scenario. These shifts were driven by earlier snowmelt and higher spring/summer temperatures. A delay in sowing up to one month may be possible because the elevated temperatures can dehydrate soil, enabling agricultural machine operation without being hindered by wet soil under RCP8.5 (Murakami *et al* 2022). While the delayed sowing dates and advanced harvesting dates imply a reduction in the growth period, this reduction appeared to be compensated by a decrease in the number of snow-covered days (figure S4). Although the durations of effective growth periods were projected to change minimally, these shifts in the phenological events may have significant effects on canopy light absorption and the balance between source and sink. The advanced growth period apart from the summer solstice may decrease canopy light absorption during the months following snowmelt. Low light availability during this period, which corresponds to the period of stem elongation to flowering, has been repeatedly linked to a decrease in grain number (i.e., sink strength), a key determinant of grain yield (e.g. Fischer 1985). On the other hand, the advanced grain-filling period, centered around the summer solstice, would increase canopy light absorption and photosynthetic carbon fixation after anthesis (i.e., source strength). These phenological shifts will therefore induce an increase in the source:sink ratio under future climate conditions. While upregulated source activity during the grain-filling period has sometimes been reported to result in an increased grain yields (e.g. Xu *et al* 2016, Shimoda and Sugikawa 2020), a decrease in sink strength is likely to have dominant and negative effects on wheat yield because sink strength typically limits wheat yield (Borrás *et al* 2004). Although several chronological analyses have suggested that the source-sink balance has shifted from strong sink limitation to co-limitation in modern cultivars (e.g. Kruk *et al* 1997, Aisawi *et al* 2015, Alvarez Prado *et al* 2023), our simulation emphasized the need to enhance sink strength through breeding and management practices. Since most crop breeding efforts for adaptation have been devoted to enhancing heat and drought tolerance (e.g. Langridge and Reynolds 2021), the present result highlights a new aspect of breeding strategies for climate change adaptation. Cooperation between meteorologists and breeders should be promoted to facilitate designing breeding goals that combine crop calendar and climate conditions.

**Figure 4:**
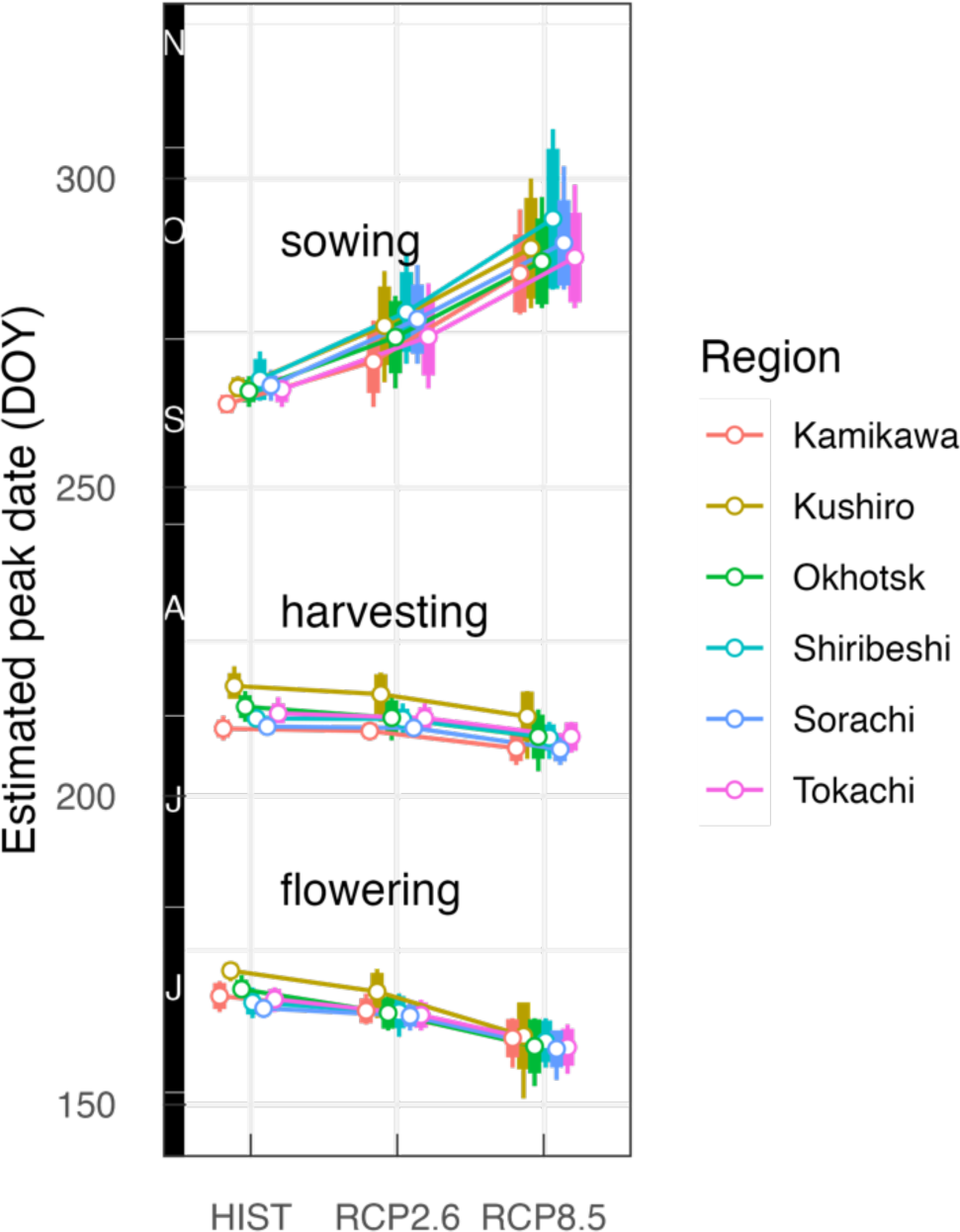
Projected shifts in peak dates (day of the year; DOY) of winter wheat phenological events in Hokkaido estimated by the phenological window model. Estimated peaks obtained under the historical climate conditions and those at the late 21st century under the RCP2.6 and RCP8.5 scenarios are shown. Thin and thick error bars represent ranges 10–90 percentiles and standard deviations, respectively.

### 2.3 Risks associated with precipitation events at unfavorable times

Analysis by two-way ANOVA showed significant effects of the two factors (RCP and subprefecture) on precipitation-associated risks around flowering and harvesting, but their interactions were not significant (table 1). The likelihood of two-day precipitation events around harvesting was projected to increase under future conditions, leading to a higher risk of preharvest sprouting under the RCP2.6 and RCP8.5 scenarios compared to HIST (table 1). This was partly because climatological precipitation around August was projected to increase in the analyzed regions (figure S5). In contrast, the number of rainy days around flowering was projected to decrease under the RCP8.5 scenario compared to the HIST and RCP2.6 scenarios (table 1). This change in precipitation patterns is in agreement with typical climate change scenarios, where the number of rainy days will decrease while total precipitation during a single rainfall event will increase (Masson-Delmotte *et al* 2021).

**Table 1:** Precipitation-associated risks at the flowering and harvesting stages under the historical climate condition (HIST) and in the late 21st century under the RCP2.6 and RCP8.5 scenarios.

| Factor | Number of rainy days around<br>flowering [d]<sup>z</sup> | Maximum two-day precipitation around<br>harvesting [mm]<sup>z</sup> |
| --- | --- | --- |
| HIST | 3.68<sup>a</sup> (2–5) | 29.6<sup>b</sup> (10.3–41.4) |
| RCP2.6 | 3.46<sup>ab</sup> (2–5) | 37.4<sup>a</sup> (11.4–52.7) |
| RCP8.5 | 3.31<sup>b</sup> (2–5) | 38.7<sup>a</sup> (9.63–54.4) |
| Climate scenario (C)<sup>y</sup> | $P = 0.0042^{**}$ | $P < 0.001^{***}$ |
| Subprefecture (S)<sup>y</sup> | $P < 0.001^{***}$ | $P < 0.001^{***}$ |
| C × S<sup>y</sup> | $P = 0.99^{\text{n.s.}}$ | $P = 0.88^{\text{n.s.}}$ |

z: Means and interquartiles of mean values for 14 subprefectures. Different letters within each column represent significantly differences among the scenarios (Tukey–Kramer’s HSD test, $P < 0.05$ ).
y: Results of two-way ANOVA (\*\*: $P < 0.01$ ; \*\*\*: $P < 0.001$ , n.s.: not significant).

To assess the transition in the opportunity loss attributed to precipitation-associated risks, we incorporated two additional rules to the window model to avoid crop failure and integrated sowing likelihood. When these risks were not considered, integrated sowing likelihood was projected to increase under future climate conditions (figure 5). However, the likelihood tended to be smaller under future climate conditions when the risks were considered in conjunction with substantial variability depending on the regions and risks considered (figure S2). This suggests that resistance and tolerance to these undesirable phenomena will be more important under future climate conditions. Because crop responses under wet conditions have been less well described and analyzed in the literature, some crop models lack representation of processes induced by wet conditions and have failed to predict yields under such conditions in wheat (e.g., van der Velde *et al* 2012, 2020) and other crops (Rosenzweig *et al* 2002, Li *et al* 2019). Developing adaptive measures and quantitative risk assessments will become increasingly important as crop production under wet conditions may become more frequent due to the projected intensification in precipitation patterns.

**Figure 5:**
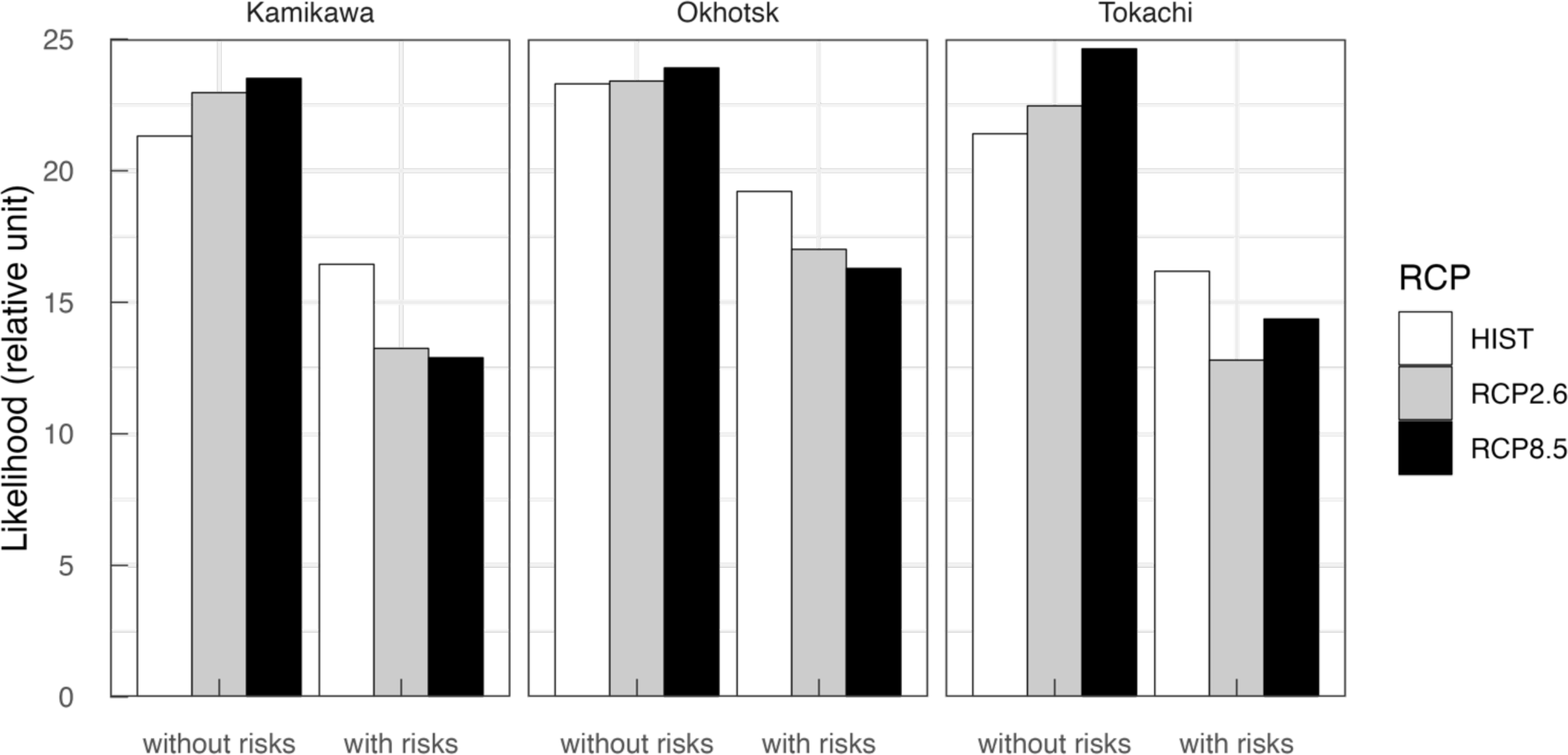
Effects of precipitation-induced crop failure on winter wheat sowing likelihoods under the historical climate condition (HIST) and in the late 21st century under the RCP2.6 and RCP8.5 scenarios. The likelihoods were calculated so that flowering and harvesting avoid pollination failure and preharvest sprouting (i.e., with risks) or without considering these risks (i.e., without risks).

Note that the present analyses of precipitation-associated risks were based on primitive assumptions. For pollination failure, most studies have focused on the effects of drought and high temperatures (e.g. Barnabás *et al* 2008). Although some studies have reported that yield loss is attributable to continuous precipitation around flowering and corresponding pollination failure (Shimoda *et al* 2022, Nóia Júnior *et al* 2023), no widely used model or index has been developed to assess this effect. While a certain amount and duration of precipitation are prerequisites for preharvest sprouting, it has been reported that sprouting only occurs when seeds are sprouting suspicious with reduced dormancy level (Mares and Mrva 2014). Seed dormancy levels are regulated by multiple environmental factors, particularly precipitation and air temperature, but also genetic variation (Mares 1993, Barnard and Smith 2009). Furthermore, it has been suggested that exposure to high temperatures followed by a rapid transition to low temperatures, referred to as cool-shock, can adversely affect flour quality even in the absence of preharvest sprouting (Mares and Mrva 2008).

## Conclusions

In this study, we presented an improved crop calendar model, which in addition to the parameters considered in the original model, incorporates the factors of vernalization and winter survival to obtain more accurate estimates of the winter wheat growing season, focusing on Hokkaido as a case study. The model simulations forced by projected climates scenarios indicated a delay in sowing up to one month accompanied by slight advances in the timing of flowering and harvesting. We also evaluated the potential risks of pollination failure and preharvest sprouting due to precipitation during key growth stages such as flowering and harvesting. The results showed that the shifts in wheat crop calendars and precipitation patterns would likely increase these risks under a warming climate. These findings are expected to be useful for setting winter wheat breeding goals and developing management practices for adaptation.

## Acknowledgements

This work was supported by JSPS KAKENHI Grant Numbers JP23K14051 (KM), JP22H00577 and JP23H00351 (TI). TI was partly supported by the Environment Research and Technology Development Fund (JPMEERF23S21120) of the Environmental Restoration and Conservation Agency provided by the Ministry of Environment.

## Ethical statement

This research did not include human subjects, human data or tissue, or animals.

## Supplemental figures

**Figure S1:**
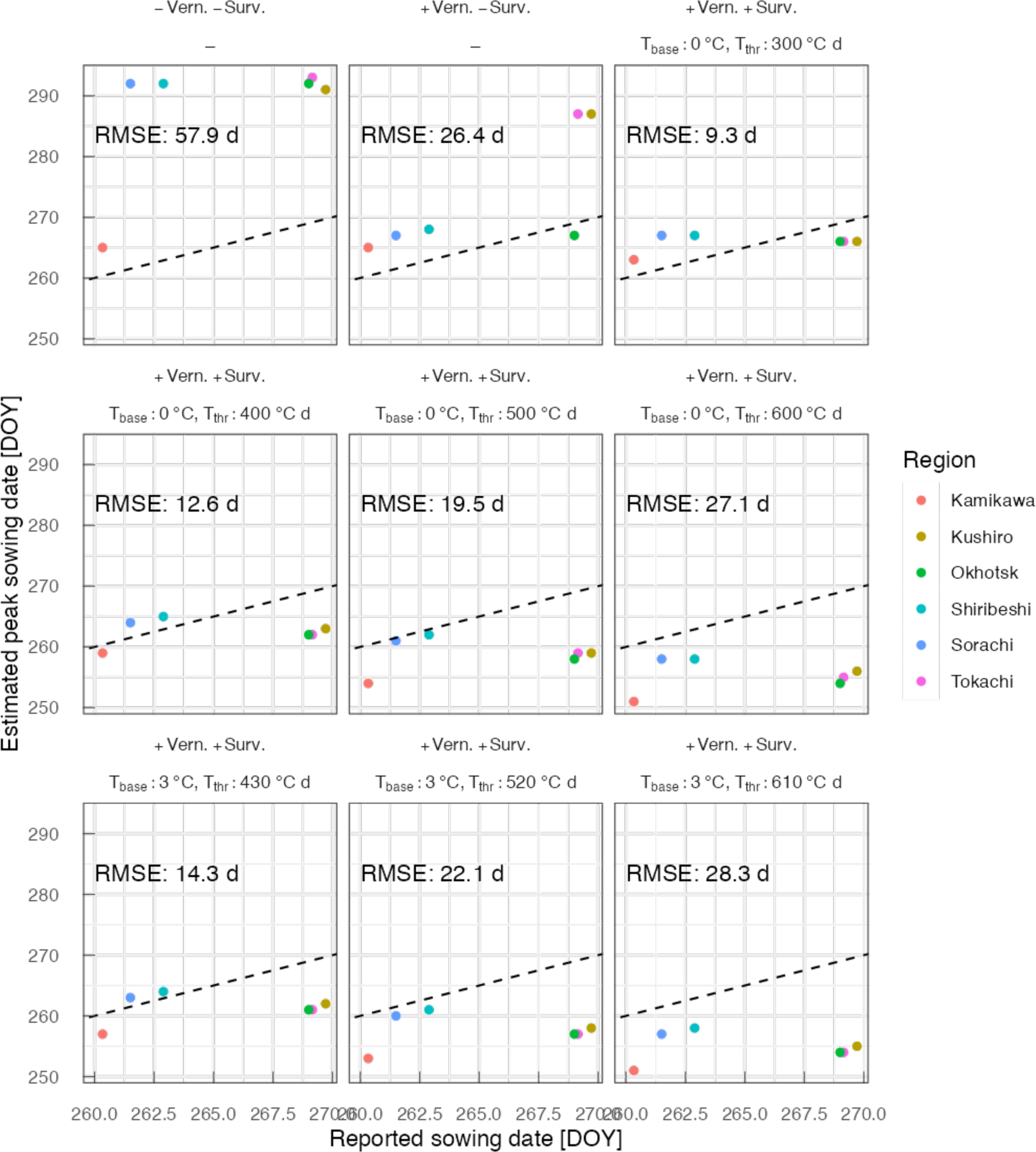
Influence of constraints of vernalization (Vern.) and winter survival (Surv.) on peak sowing dates estimated by the phenological window model. Dates estimated by models with (+) or without (-) these constraints, as well as several combinations of parameters necessary for evaluating winter survival, are compared to those reported by the government office of Hokkaido (2000–2020). Winter survival was considered to be successful if the cumulative effective temperature with a base temperature (Tbase) from the sowing date to the commencement of the long-term snow period was greater than a threshold value (Tthr). DOY: Day of year. RMSE: Root mean square error.

**Figure S2:**
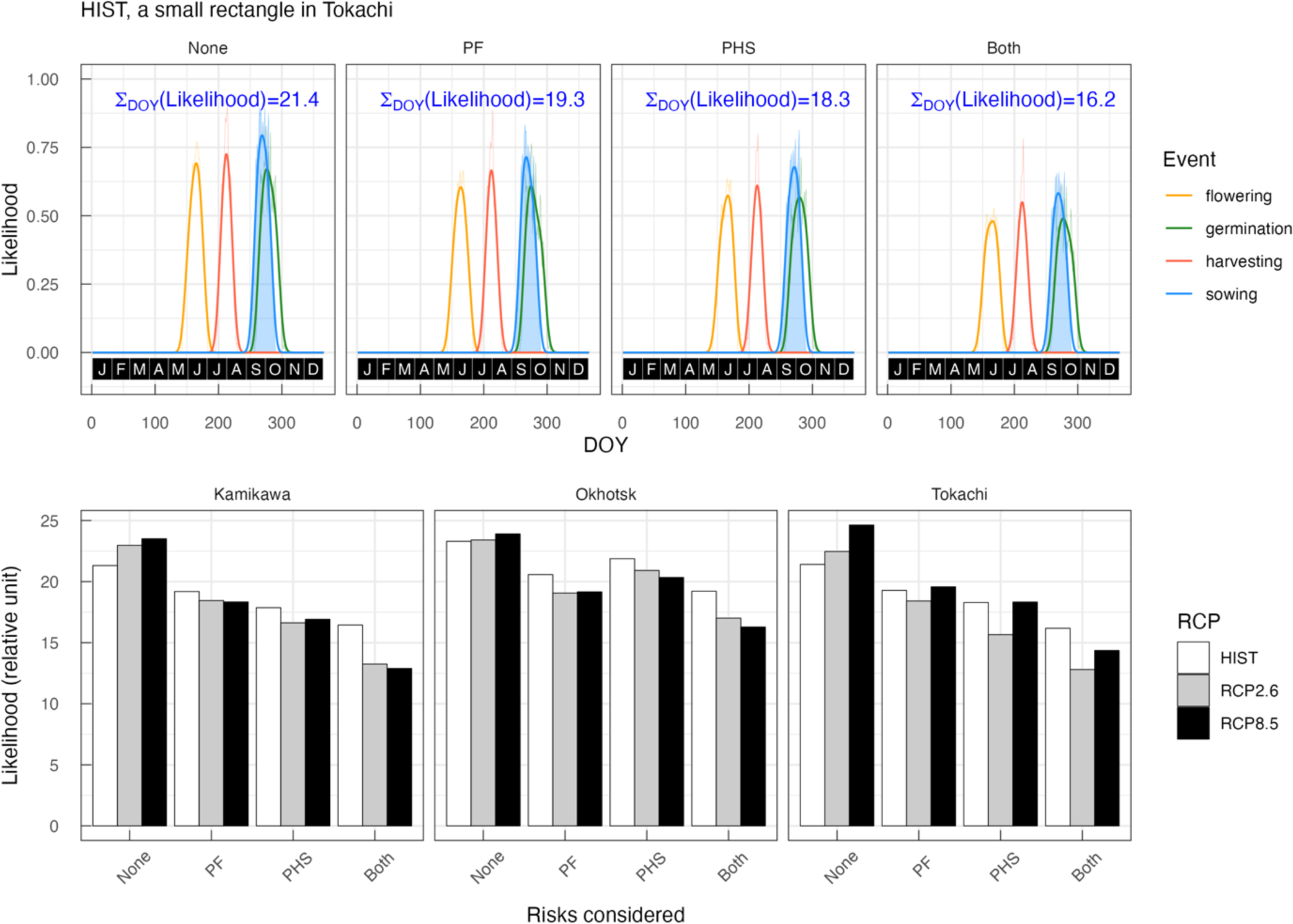
Estimates of sowing likelihood of precipitation-associated opportunity loss of winter wheat cultivation (top panels). Phenological event windows at a given site and climate are simulated with or without considering risks of pollination failure (PF) and/or preharvest sprouting (PHS). Daily likelihood values are integrated over time (i.e., blue areas). Effects of PF and PHS on sowing likelihoods under the historical climate condition (HIST) and in the late 21st century under the RCP2.6 and RCP8.5 scenarios (bottom panels). Sowing likelihoods estimated for three small regions in different subprefectures (square insets in figure S1) are shown.

**Figure S3:**
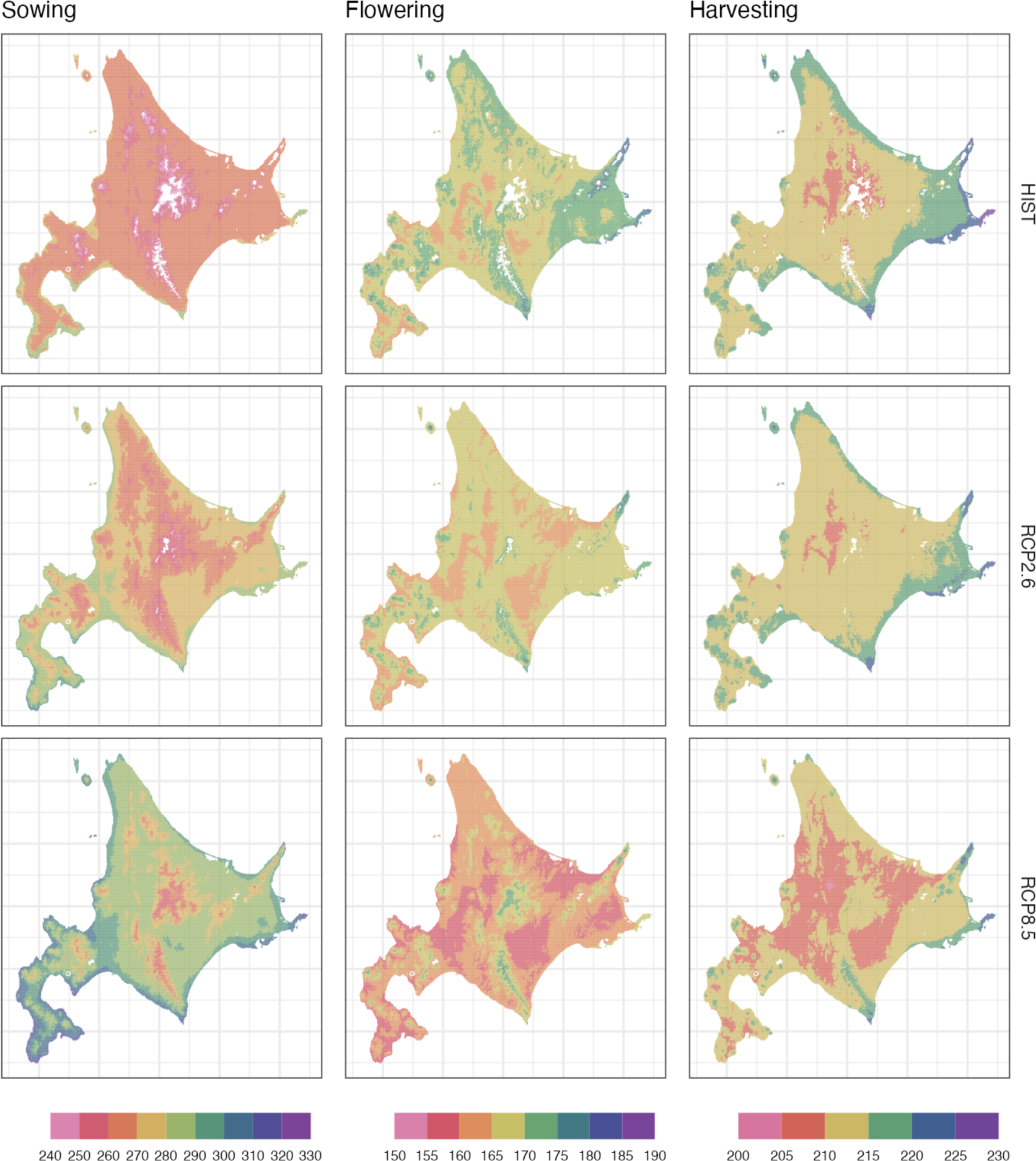
Projected peak dates (DOY) of sowing, flowering, and harvesting of winter wheat the historical climate condition (HIST) and in the late 21st century under the RCP2.6 and RCP8.5 scenarios.

**Figure S4:**
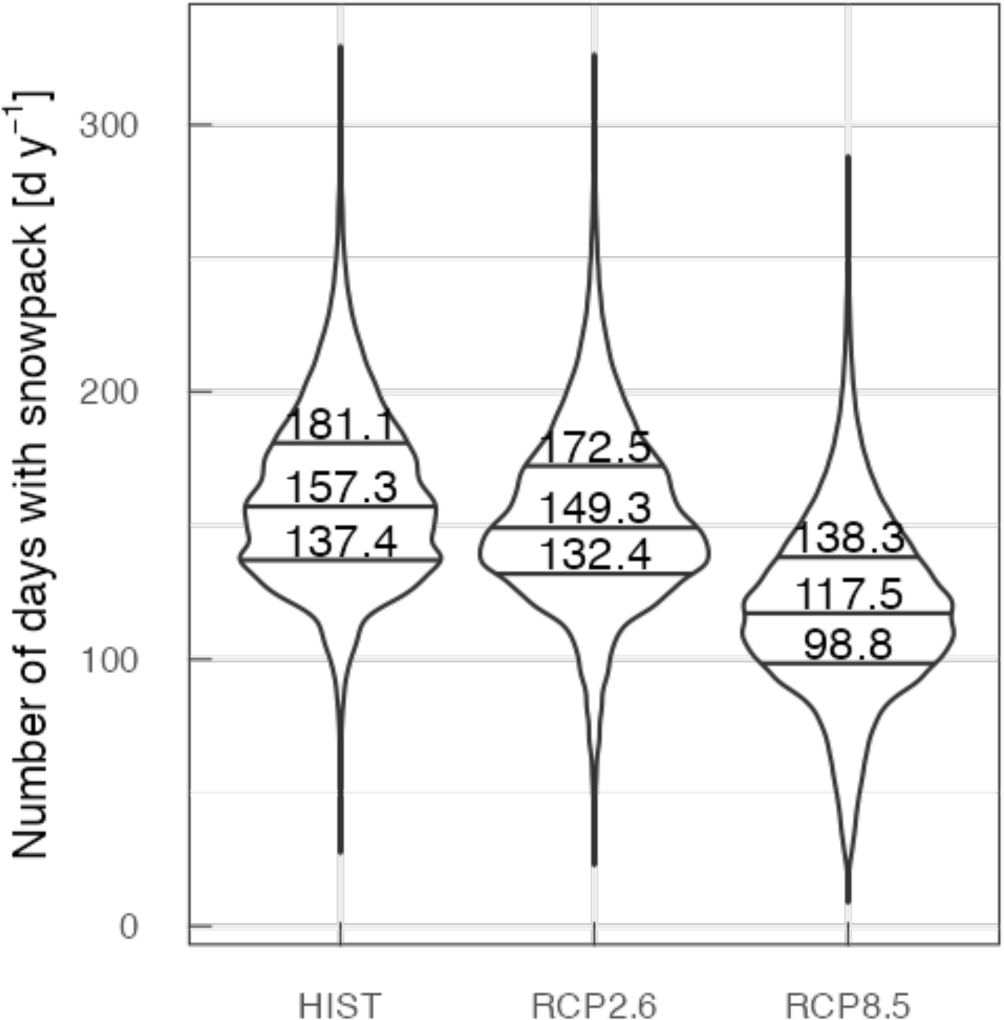
Number of snow-covered days in Hokkaido under the historical climate condition (HIST) and in the late 21st century under the RCP2.6 and RCP8.5 scenarios. Medians and interquantiles of all grid cells over Hokkaido are shown.

**Figure S5:**
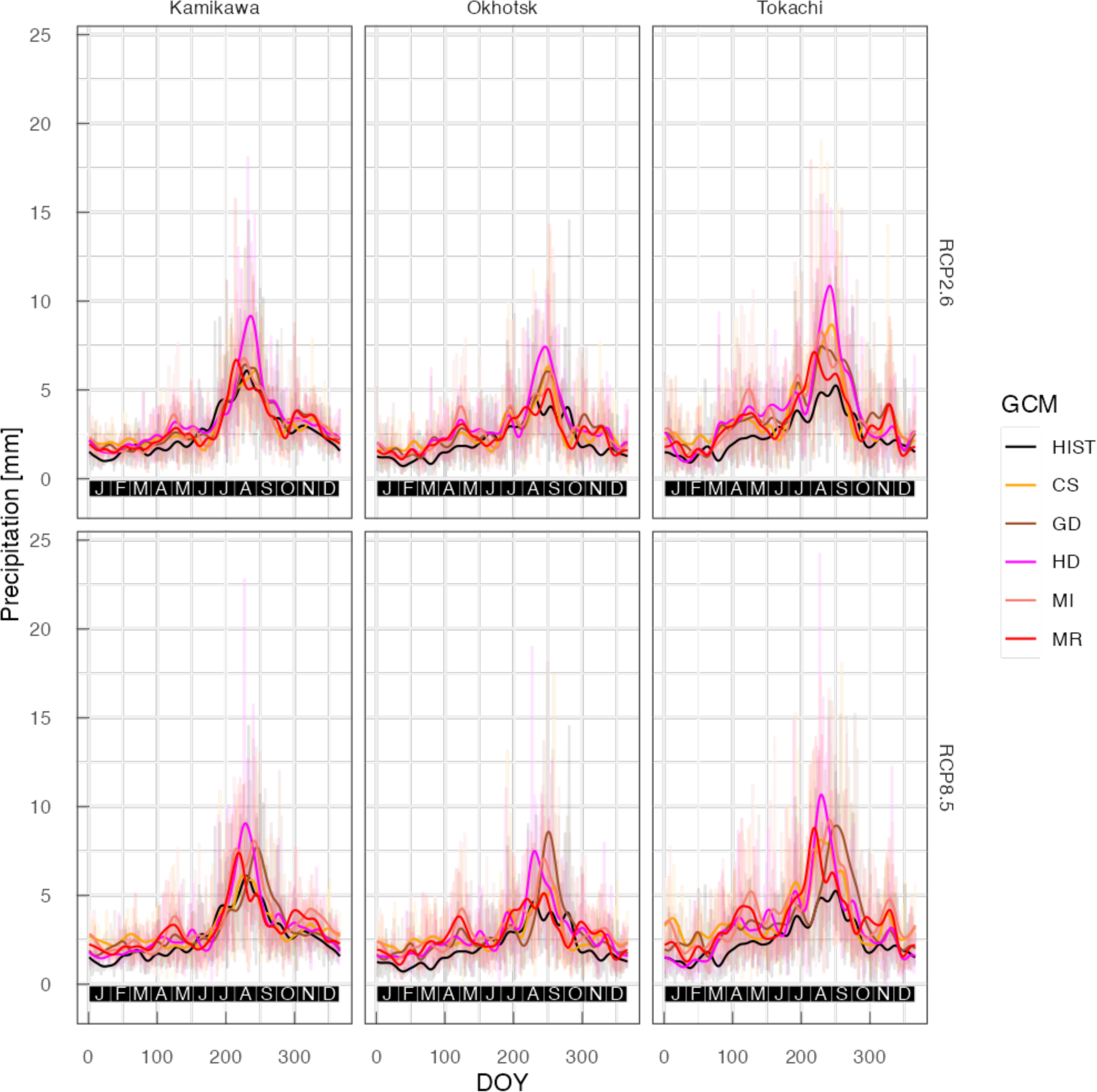
Climatological daily precipitation in three regions (square insets in figure 1) under the historical climate condition (HIST; 2000–2020) and in the late 21st century under the RCP2.6 and RCP8.5 scenarios (2070–2090) derived from five global climate models (GCMs). Thin and thick lines indicate the raw model output and smoothed data (7-day moving average repeated 10 times), respectively. CS: CSIRO-Mk3.6.0, GD: GFDL-CM3, HD: HadGEM2-ES, MI: MIROC5, MR: MRI-CGCM3.

